# Checkpoint phosphorylation sites on budding yeast Rif1 protect nascent DNA from degradation by Sgs1-Dna2

**DOI:** 10.1101/2020.06.25.170571

**Authors:** Chandre Monerawela, Shin-ichiro Hiraga, Anne D. Donaldson

**Affiliations:** Institute of Medical Sciences, University of Aberdeen, Aberdeen AB25 2ZD, Scotland, UK

## Abstract

In budding yeast the Rif1 protein is important for protecting nascent DNA at blocked replication forks, but the mechanism has been unclear. Here we show that budding yeast Rif1 must interact with Protein Phosphatase 1 to protect nascent DNA. In the absence of Rif1, removal of either Dna2 or Sgs1 prevents nascent DNA degradation, implying that Rif1 protects nascent DNA by targeting Protein Phosphatase 1 to oppose degradation by the Sgs1-Dna2 nuclease-helicase complex. This functional role for Rif1 is conserved from yeast to human cells. Yeast Rif1 was previously identified as a target of phosphorylation by the Tel1/Mec1 checkpoint kinases, but the importance of this phosphorylation has been unclear. We find that nascent DNA protection depends on a cluster of Tel1/Mec1 consensus phosphorylation sites in the Rif1 protein sequence, indicating that the intra-S phase checkpoint acts to protect nascent DNA through Rif1 phosphorylation. Our observations uncover the pathway by which budding yeast Rif1 stabilises newly synthesised DNA, highlighting the crucial role Rif1 plays in maintaining genome stability from lower eukaryotes to humans.

**Author summary:** Genome instability is a leading factor contributing to cancer. Maintaining efficient error-free replication of the genome is key to preventing genome instability. During DNA replication, replication forks can be stalled by external and intrinsic obstacles, leading to processing of nascent DNA ends to enable replication restart. However, the nascent DNA must be protected from excessive processing to prevent terminal fork arrest, which could potentially lead to more serious consequences including failure to replicate some genome sequences. Using a nascent DNA protection assay we have investigated the role of the budding yeast Rif1 protein at blocked replication forks. We find that Rif1 protects nascent DNA through a mechanism that appears conserved from yeast to humans. We show that budding yeast Rif1 protects nascent DNA by targeting Protein Phosphatase 1 activity to prevent degradation of nascent DNA by the Sgs1-Dna2 helicase-nuclease complex. Furthermore, we find that Rif1 phosphorylation by the checkpoint pathway during replication stress is crucial for this function. Our results indicate that the S phase checkpoint machinery acts by phosphorylating Rif1 to protect nascent DNA, providing important clues concerning the conserved role of Rif1 in regulating events when replication is challenged.

## Introduction

Maintaining genome integrity during replication of the genome is key to preventing oncogenesis. During S phase of the cell cycle, when DNA replication occurs, replication forks can encounter many obstacles that challenge error-free duplication of the genome. Numerous cellular proteins act to ensure the complete and accurate transmission of genomic information to daughter cells in each cell cycle.

Rif1 is one such protein, important for maintaining genome integrity at several steps of the chromosome cycle. Rif1 is a multi-functional protein conserved from budding yeast to humans, which was originally identified as a negative regulator of telomere length in budding yeast *Saccharomyces cerevisiae* [1]. While its telomere length regulation function appears to be specific to yeast [2], other roles of Rif1 are conserved in eukaryotes [3]. One apparently conserved function of Rif1 is promotion of double-stranded break (DSB) repair through non-homologous end joining (NHEJ). Rif1 drives DSB repair toward NHEJ by protecting 5’ ends from resection that would favour homology-directed repair (HDR), in a function that appears to be conserved from budding yeast to human cells [4–7].

Mammalian Rif1 also plays a role in programmed genomic rearrangements in mammalian cells, such as immunoglobulin class switching, which is a specialised form of NHEJ [4,5]. Another conserved function of Rif1 is control of the initiation of DNA replication [8–10]. In controlling DNA replication, Rif1 acts by suppressing premature activation of the minichromosome maintenance protein (MCM) complex as the replicative helicase. In this role Rif1 operates as a substrate-targeting subunit for Protein Phosphatase 1 (PP1), directing dephosphorylation of the MCM complex, and counteracting its phosphorylation by the Dbf4-dependent kinase (DDK) to constrain replication origin activation [11–16].

Rif1 also functions at later stages of the DNA replication process. In particular, it was recently demonstrated that mammalian Rif1 protects nascent DNA at replication forks challenged by replication inhibitors [17,18]. DNA replication forks can be impeded or stalled for many reasons. Obstacles such as collisions between the replication and transcription machinery, DNA/RNA hybrids (R-loops), ribonucleotide incorporation, DNA lesions and adducts, DNA secondary structure, repetitive DNA sequences, non-histone protein-DNA complexes, and accumulation of topological stresses may cause replication forks to stall or collapse [19,20]. Stalled forks are frequently processed to prepare them for replication restart, with the nascent DNA subject to controlled degradation to create a single-stranded stretch that can be utilised for homology-dependent fork restart mechanisms [21]. In this context degradation can be carried out by multiple nucleases, including MRN, EXO1, and DNA2 [22,23]. The action of these nucleases is restricted by a number of different proteins. In mammalian cells, BRCA1 and BRCA2 protect nascent DNA from degradation by MRE11 nuclease [24], whereas BOD1L protects the nascent DNA from the DNA2-WRN nuclease-helicase complex but not from MRE11 [25]. Human Rif1 was shown to protect the nascent DNA specifically from degradation by DNA2-WRN nuclease-helicase complex, in a function that depends on Rif1 interaction with PP1. Phosphorylation of DNA2 and WRN was increased in cells depleted for Rif1, suggesting that Rif1-PP1 could potentially modulate the phosphorylation status of DNA2-WRN to control its activity [17,18].

In budding yeast, the proteins that process stalled replication forks are less well understood. While genetic studies indicate that a similar set of proteins as in human cells are important to protect cells from replication stress, their precise molecular roles remain unclear [26–29]. It was recently reported that the budding yeast MRX protein complex (composed of Mre11, Rad50, and Xrs2) promotes resection at stalled forks, but MRX appears to act in this role by supporting the remodelling of nascent chromatin, rather than through its nuclease activity [26]. However, differences in the methodologies used make it difficult to precisely align results obtained from yeast with our knowledge of mammalian cell pathways. Based on results obtained from a fibre labelling assay similar to that typically used in mammalian cells, we recently reported that yeast Rif1 protein plays a role in protecting nascent DNA from degradation when replication is inhibited by hydroxyurea (HU). This role for yeast Rif1 is correlated with its recruitment to stalled forks [30]. The discovery aligns with the findings that mouse and human Rif1 function in nascent DNA protection [17,18], and opens the possibility of investigating the process using yeast molecular genetic tools.

Throughout eukaryotes, inhibition of DNA replication causes activation of the replication, or intra-S phase checkpoint machinery. In studying how cells respond to replication stress to maintain genome integrity, HU has been used extensively as a model drug. HU acts by inhibiting ribonucleotide reductase leading to depletion of cellular deoxyribonucleotide triphosphate (dNTP) pools, slowing down the progression of active replication forks in the cell [31]. Inhibition of DNA synthesis generates increased single-stranded DNA as the replicative helicase proceeds uncoupled from DNA synthesis [32], causing activation of the intra-S phase checkpoint through recognition of Replication Protein A (RPA) bound to single-stranded DNA (ssDNA) [19,33]. The Mec1 apical kinase is recruited to the RPA-coated ssDNA, and through the mediator kinase Mrc1 activates the effector kinase Rad53 [34]. This results in a cascade of cellular responses, mediated through phosphorylation of multiple factors by Mec1 and Rad53 [33]. Activation of the intra-S phase checkpoint strongly affects the activation of further replication origins. Globally, new origin initiation events are inhibited, but in the proximity of stalled forks dormant origins are activated, at least in mammalian cells [35,36]. At stalled replication forks the intra-S phase checkpoint is proposed to stabilise the replisome structure [37]. However, any implication of the S phase checkpoint in controlling the resection of nascent DNA at stalled forks has remained unclear.

Here we examine the mechanism by which budding yeast Rif1 protects nascent DNA at stalled forks. Using a nascent DNA protection assay we find that Rif1-PP1/Glc7 interaction is crucial to protect newly synthesized DNA during a HU block, operating to protect against Sgs1-Dna2 mediated degradation. We also reveal that a cluster of Tel1/Mec1 phosphorylation sites in the C-terminal half of the Rif1 protein sequence is important to protect nascent replicated DNA at stalled forks, indicating that the replication checkpoint machinery stimulates protection of newly-replicated DNA by phosphorylating Rif1.

## Results

### Rif1 interaction with PP1/Glc7 is crucial to protect nascent DNA from degradation under replication stress

Deletion of budding yeast Rif1 was reported to cause a defect in the protection of nascent DNA from degradation when replication fork progression is blocked by HU treatment [30]. To investigate this function of Rif1 further, we used a previously established DNA combing assay to assess the stability of labelled nascent DNA in budding yeast [30]. Briefly, cells capable of incorporating thymidine analogs were synchronized in G1 phase, then released to begin DNA replication in medium containing 5-iodo-2’-deoxyuridine (IdU). After 18 minutes, IdU was removed and HU was added, and samples collected at time points thereafter (Fig 1A). DNA from these samples was combed, and IdU-labelled nascent DNA tracts visualized by immunodetection (Fig 1B). Stability of the nascent DNA tracts during the replication block can be assessed by comparing the lengths of the nascent DNA tracts when HU is added with the tract lengths at later time points.

**Fig 1.**
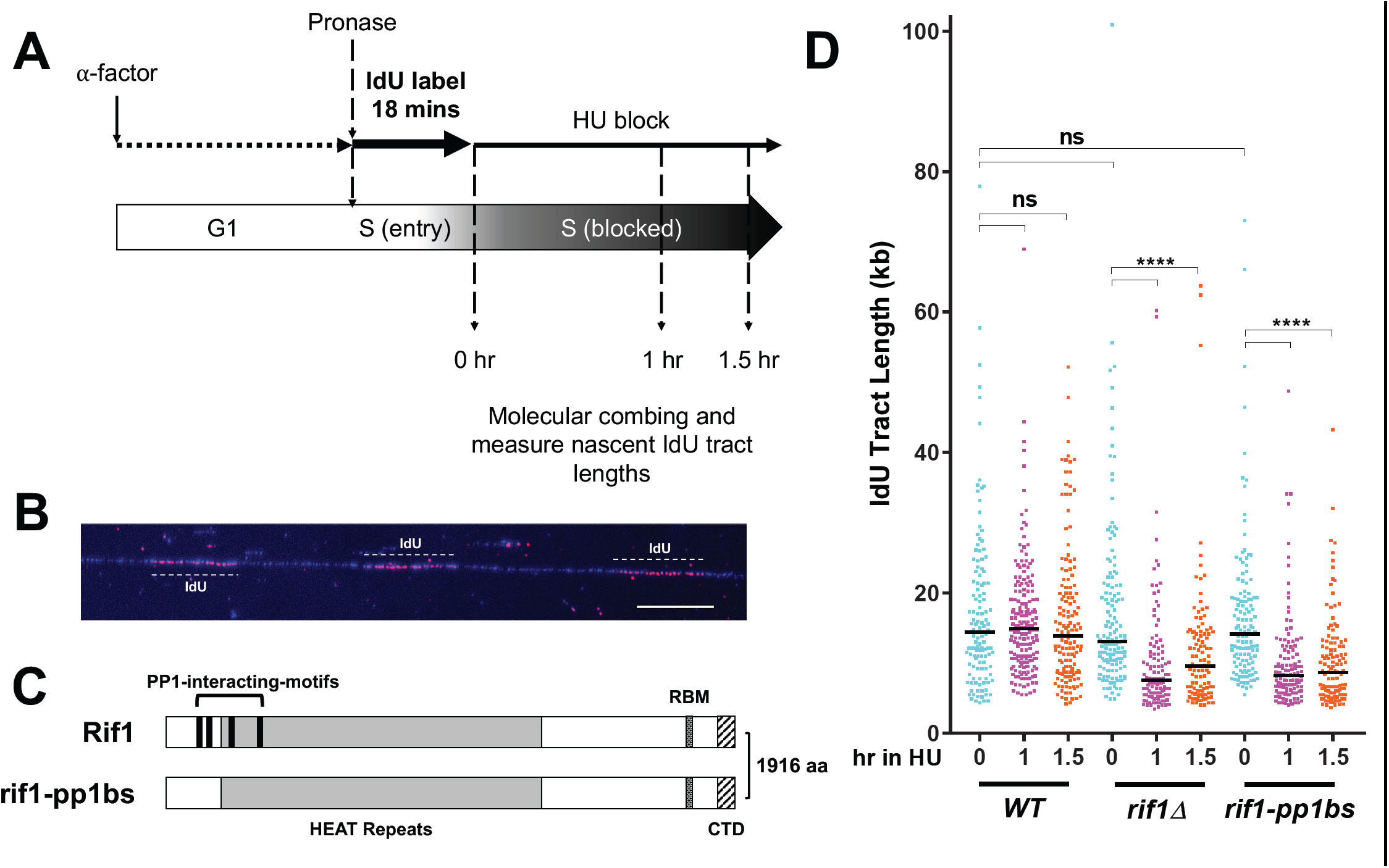
Rif1 must interact with Protein Phosphatase 1 (Glc7) to prevent degradation of nascent DNA. (A) Nascent DNA protection assay procedure. Cells arrested with α-factor were released into medium containing 1.13 mM IdU to label nascent DNA. After labelling nascent DNA for 18 mins, cells were collected by filtration and resuspended in medium containing 0.2 M HU. (B) Specimen fibre with DNA (stained blue) showing three IdU tracts (pink). Scale bar 10 µm, equivalent to 20 kb. (C) Schematic of Rif1 showing the four PP1/Glc7-interaction motifs mutated in *rif1-pp1bs*, a series of HEAT repeats, Rap1-binding motif (RBM) and C-terminal domain (CTD), a low-affinity Rap1 binding site and tetramerization module [6]. (D) Degradation of nascent DNA in *rif1-pp1bs* cells. Plot shows distribution of IdU tract lengths after incubation in a HU block for the intervals specified. ≥100 tracts were measured for each condition. In this and subsequent plots, black horizontal bars indicate median values, **** indicates *p*-values less than 0.0001 and were obtained by Mann-Whitney-Wilcoxon test, ns means “not significant”. Strains used were VGY85 (*WT*), CMY6 (*rif1Δ*) and CMY42 (*rif1-pp1bs*)

Association with PP1 is essential for various Rif1 functions in budding yeast and humans [11,12,15,17,38]. In mammalian cells nascent DNA protection by Rif1 requires PP1 interaction, so we tested if Rif1-PP1/Glc7 interaction is also needed to protect nascent DNA in budding yeast [17]. For this purpose, we used a *RIF1* allele (*rif1-pp1bs*) that has all four PP1-interacting motifs mutated (Fig 1C) [12]. Nascent DNA tract length labelled during the 18 min incubation did not significantly differ between wild type (*WT*), *rif1Δ* and *rif1-pp1bs* cells, revealing that the distance travelled by forks in this interval was similar in all three strains at the time of HU addition (0 hr, Fig 1A & D). However while nascent DNA tract length in *WT* cells did not change during the HU treatment, the median length of *rif1Δ* nascent DNA tracts was significantly decreased by 1 hr into the HU block (from 13.8 kb to 7.5 kb, Fig 1D) confirming the previous finding that Rif1 is required to prevent degradation of nascent DNA in *S. cerevisiae* [30]. In *rif1-pp1bs*, after 60 minutes in the HU block, we saw a similar decrease in the length of newly synthesized DNA tracts (from 14.5 kb to 8.2kb; Fig 1D). These results indicate that Rif1 must interact with PP1 to prevent degradation of nascent DNA under replication stress in budding yeast, as in human cells.

### Sgs1, the yeast WRN helicase homolog, mediates nascent DNA degradation when protection by Rif1 is lacking

In budding yeast several nucleases and helicases have been implicated in resecting nascent DNA ends [26] after remodelling by the MRX complex. Studies in mammalian cells also demonstrated that several different proteins protect nascent DNA from degradation by specific exonucleases [25,39–42]. We tested which is the major nuclease and/or helicase responsible for nascent DNA resection in budding yeast lacking Rif1 function. We first examined the effect of deleting the 5’ to 3’ Exo1 nuclease, as in Fig 1A. If Rif1 is in charge of protecting the nascent DNA from degradation by Exo1, deletion of *EXO1* should rescue the nascent DNA degradation phenotype of *rif1Δ*. Removal of Exo1, however, did not significantly affect the initial replicated tract length after the 18 min IdU labelling period, either in the *RIF1* (*WT*) or the *rif1Δ* background (Fig 2A). After 1 hr and 1.5 hr of HU blocking, the *rif1Δ exo1Δ* mutant showed a significant decrease in IdU-labeled tract length (from 11.0 kb to 8.4kb after 1 hour HU treatment; Fig 2A), similar to the decrease of 12.4 kb to 7.2 kb observed in the *rif1Δ* single mutant. Therefore nascent DNA degradation still occurs when Exo1 is absent, indicating that Exo1 is not the major nuclease responsible for degrading nascent DNA when Rif1 function is lacking.

**Fig 2.**
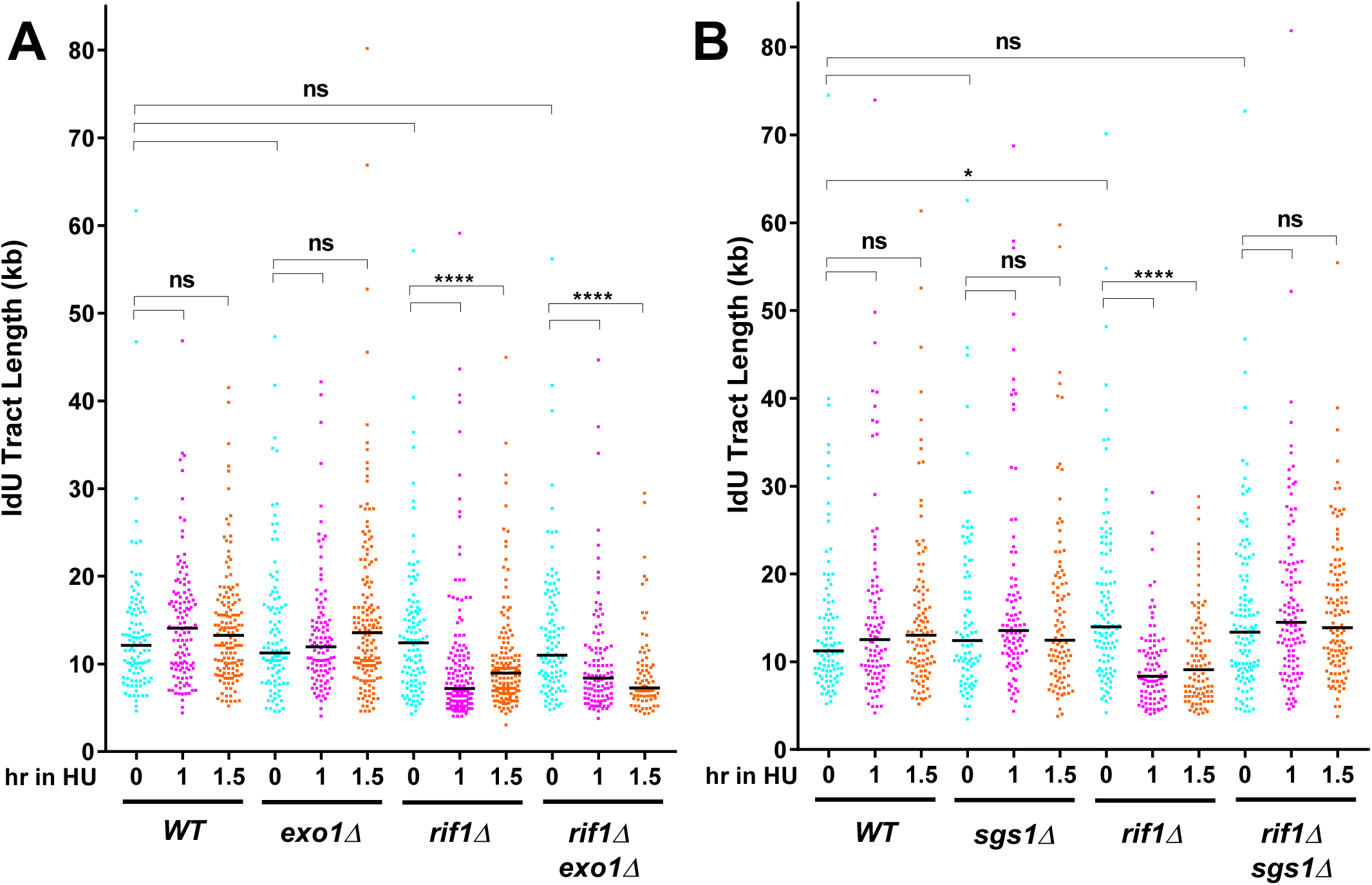
Sgs1 mediates degradation of nascent DNA in *rif1Δ* mutants. (A) *EXO1* deletion does not prevent the degradation of nascent DNA in a *rif1Δ* mutant. (B) *SGS1* deletion does prevent the degradation of nascent DNA in a *rif1Δ* mutant. IdU tracts were labelled and lengths analysed as in Fig 1. Strains used were VGY85 (*WT*), CMY6 (*rif1Δ*), CMY46 (*exo1Δ*), CMY47 (*rif1Δ exo1Δ*), CMY52 (*sgs1Δ*) and CMY53 (*rif1Δ sgs1Δ*).

We next examined whether Sgs1, the yeast homolog of the human WRN, is important for nascent DNA degradation. Results from *sgs1Δ* and *rif1Δ sgs1Δ* strains show no significant difference in the median length of newly synthesized DNA tracts when compared to *WT* and *rif1Δ*, respectively (Fig 2B), indicating that removal of Sgs1 does not affect the progression of unblocked replication forks in the initial labelling period. However, when compared to the strong resection phenotype of *rif1Δ* cells after HU addition, *rif1Δ sgs1Δ* cells showed no significant decrease in nascent DNA tract length even after 1.5hr HU treatment (Fig 2B). Therefore, deleting Sgs1 rescues the degradation phenotype seen in a *rif1Δ* background, indicating that the budding yeast Sgs1 helicase is important for degrading nascent DNA deprotected by Rif1 removal. These observations are altogether consistent with findings in human cells lacking Rif1, where the WRN helicase was needed for nascent DNA degradation [17].

### Dna2 is required for the degradation of nascent DNA in cells lacking Rif1

Sgs1 acts as a helicase that unwinds dsDNA to feed an ssDNA strand to the nuclease Dna2, promoting processive resection of DNA ends [43]. Since nascent DNA degradation in the absence of yeast Rif1 requires Sgs1, we explored whether Dna2 is the major nuclease activity responsible for degradation of newly synthesized DNA in a *rif1Δ* background.

Dna2 is an essential protein, probably because of its involvement in Okazaki fragment processing [44]. We first investigated a temperature sensitive *dna2-1* allele and investigating the effect of combining it with *rif1Δ*, however, the strain background was unsuitable for the assay due to poor growth even at its permissive temperature [45] (Fig S1). Therefore, we instead designed an inducible degradation strategy to test the involvement of Dna2 in nascent DNA degradation.

Henceforth, we adopted an auxin-inducible degradation (AID) system [46] to enable inducible degradation of Dna2 protein, only in a desired time period during experiments. An AID tag was fused to the C-terminus of Dna2 in *WT* and *rif1Δ* cells, in a strain background bearing a cassette encoding OsTIR1, the E3 ubiquitin ligase that promotes degradation of AID-tagged proteins [47]. OsTIR1 is under the control of a Galactose-regulated promoter, enabling induced depletion of Dna2-AID by addition of Galactose and auxin. We confirmed that cells with Dna2-AID were unable to grow on plates containing galactose (YP-Gal) and auxin (Fig 3A). Western blot analysis confirmed that in α-factor-blocked cultures, Dna2-AID was swiftly degraded and became undetectable 15 min after auxin addition (Fig 3B).

**Fig 3.**
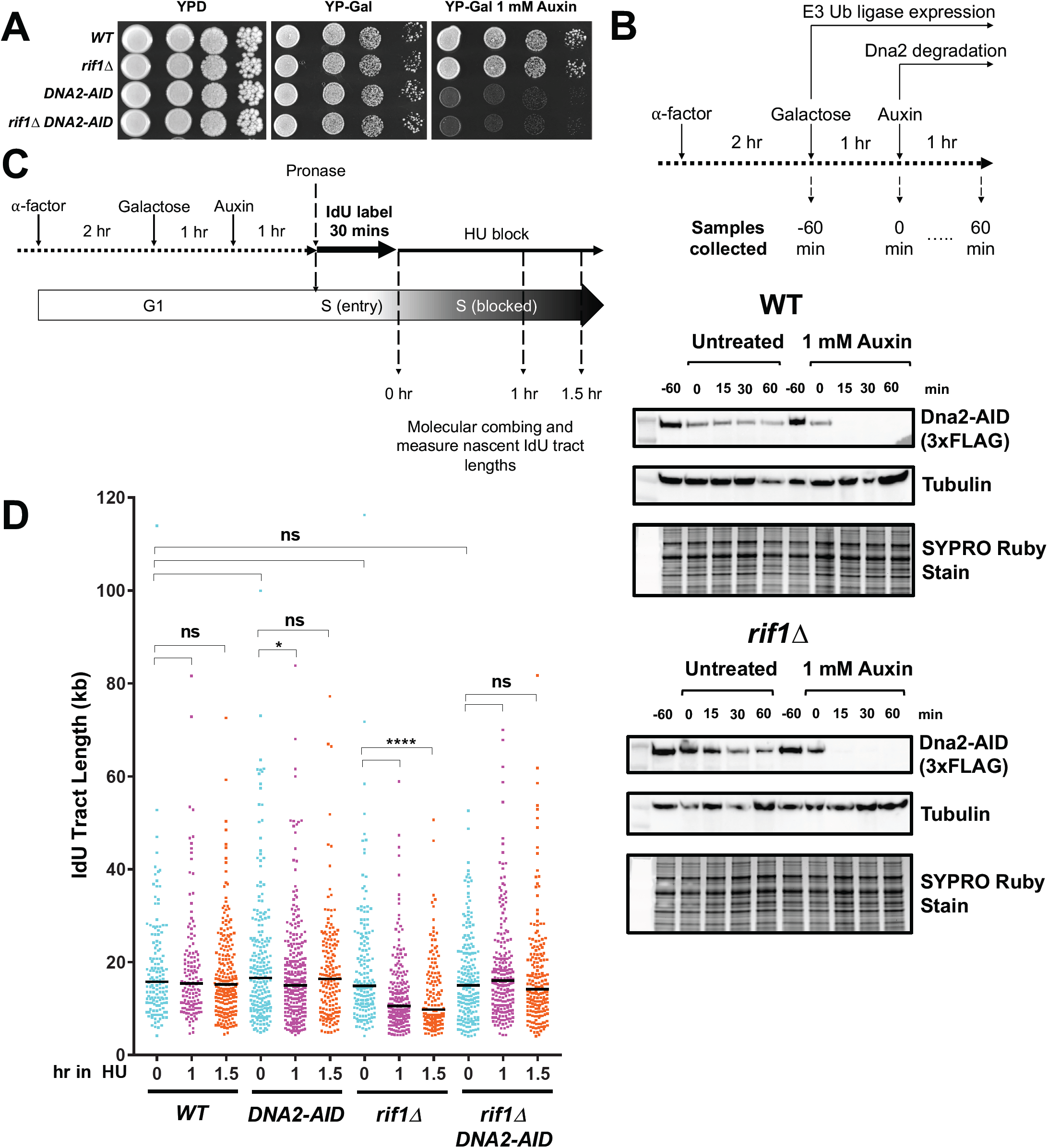
Rif1 protects nascent DNA from Dna2 mediated degradation. (A) Dna2 depletion via an AID degron tag prevents cell growth. Serial dilutions (1:10) of cells were grown on YPD, YP-Gal and YP-Gal supplemented with 1 mM auxin at 30°C. (B) Western blot using anti-FLAG antibody confirms degradation of Dna2 in wild type (middle panel) and *rif1Δ* (lower panel) cells. As illustrated in the schematic (top), samples were collected from G1-arrested cultures 60 mins before auxin addition, and then at a series of timepoints after auxin addition. (C) Nascent DNA protection assay procedure to test effects of Dna2 depletion, incorporating OsTIR1 induction by galactose and auxin addition to degrade Dna2. (D) Depletion of Dna2-AID prevents degradation of nascent DNA in *rif1Δ* cells. Here and below * and **** indicate *p*-values less than 0.05 and 0.0001, respectively. Strains used were CMY54 (*WT*), CMY56 (*rif1Δ*), CMY58 (*DNA2-AID*) and CMY59 (*rif1Δ DNA2-AID*).

To test nascent DNA protection using Dna2-AID tagged cells, we modified the experimental procedure as shown in Fig 3C. Cells were first synchronized in G1 phase, then OsTIR1 was induced by addition of galactose. 1 hr later auxin was added to deplete Dna2. Replication proceeds somewhat more slowly in galactose than in glucose medium, so the initial IdU labelling period was extended to 30 min, prior to addition of HU to block replication (Fig 3C). Samples were then taken 0, 1 and 1.5 hr after HU addition for nascent DNA combing analysis.

Removal of Dna2 did not impact the initial synthesis of DNA during the 30 min IdU labelling, in either a *WT* or *rif1Δ* background (Fig 3D, compare 0 hr samples). The nascent DNA protection defect of *rif1Δ* cells was still apparent using this modified procuedure (Fig 3D, *rif1Δ* cells 1 and 1.5 hr samples), with the median IdU-labelled tract length significantly decreased (from 14.9 kb to 9.8 kb) after 1.5 hours in the HU block. Depleting Dna2-AID in the *rif1Δ* background however largely prevented the degradation phenotype, with only a slight decrease in nascent DNA tract lengths over the course of the 1.5 hr HU block, which was not statistically significant (Fig 3D, *rif1Δ DNA2-AID*, 1 and 1.5 hr samples). We conclude that Dna2 is the major nuclease responsible for degradation of nascent DNA in *rif1Δ* cells. In human cells also, nascent DNA deprotected by loss of Rif1 is degraded primarily by Dna2 [17,18]. Therefore, our results confirm that Rif1 protects nascent DNA from degradation by the Dna2-WRN/Sgs1 nuclease-helicase in yeast as well as in human cells, in the pathway that appears to be evolutionarily conserved.

### Phoshorylation of a cluster of S/TQ checkpoint recognition sites in yeast Rif1 is required for replication fork protection

The Rif1 sequence contains a cluster of seven ‘S/TQ’ recognition motifs for the PIKK checkpoint kinases Tel1 and Mec1, located between amino acids 1308 and 1570 (Fig. 4A). This region has been termed the ‘SCD’ (for ST/Q Cluster Domain) [48]. Four of these potential phosphorylation sites have been confirmed to be phosphorylated *in vivo* [49] by mass spectrometry analysis of immuno-precipitated Rif1 from yeast cells. Phosphorylation of two of these sites depends on Tel1 or Mec1 [48–50], suggesting some aspect of Rif1 function is controlled by the checkpoint machinery. However, the functional importance of these checkpoint phosphorylation sites has remained unclear.

**Fig 4.**
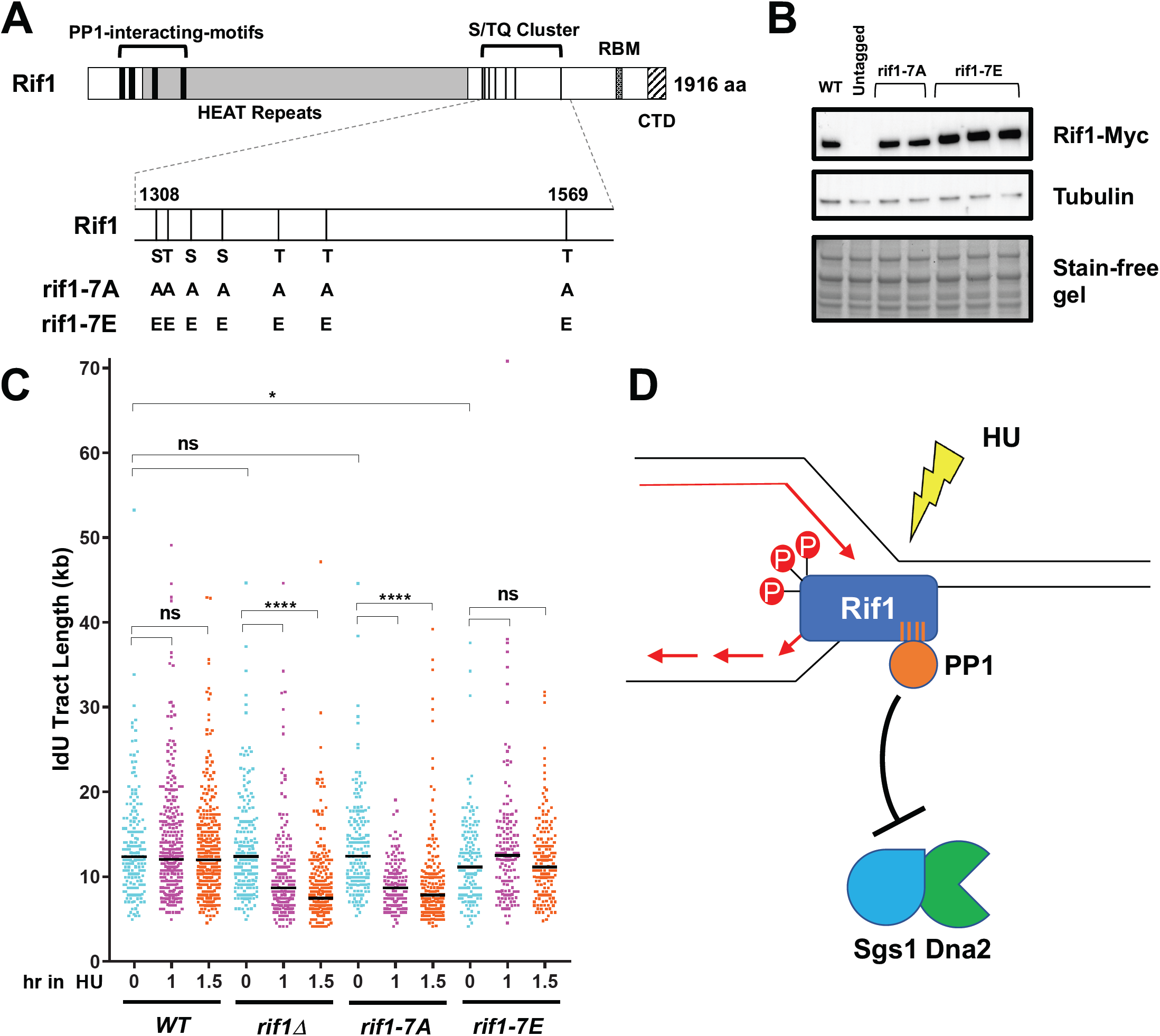
Nascent DNA protection requires a cluster of Tel1/Mec1 checkpoint recognition sites in the unstructured Rif1 C-terminal domain. (A) A schematic illustration of Rif1 structure including seven S/TQ sites mutated to alanines in the phospho-dead mutant (*rif1-7A*) or glutamic acids in the phosphomimic mutant (*rif1-7E*). (B) Western blot testing expression of *rif1-7A* and *rif1-7E* alleles. Strains were YSM20 (WT), YK402 (untagged), CMY135-6 (*rif1-7A*) and CMY137-9 (*rif1-7E*). (C) Phospho-dead mutant *rif1-7A* shows a profound defect in the protection of nascent DNA at HU-blocked forks, while nascent DNA protection is intact in the *rif1-7E* phospho-mimic mutant. Black horizontal bars indicate median values. Strains used were VGY85 (*WT*), CMY6 (*rif1Δ*), CMY130 (*rif1-7A*) and CMY132 (*rif1-7E*). (D) Model for protection of nascent DNA by *S. cerevisiae* Rif1 under HU-induced replication stress. Replication fork stalling upon HU treatment causes activation of the intra-S phase checkpoint, and consequent phosphorylation of the S/TQ site cluster in Rif1 (red circle), enabling Rif1-PP1 at the stalled replication fork to oppose degradation by the Sgs1-Dna2 complex, potentially through direct dephosphorylation.

It was previously demonstrated that *rif1-ΔC594*, a C-terminal truncation mutant of Rif1, is defective for protection of nascent DNA after HU treatment [30]. The Rif1-ΔC594 protein ends at amino acid 1322 and lacks five of the seven clustered consensus checkpoint phosphorylation sites in the S/TQ site cluster, as well as the C-terminal telomere interaction domain, raising the possibility that checkpoint phosphorylation contributes to nascent DNA protection by Rif1. We investigated whether phosphorylation within the S/TQ site cluster is important for Rif1 to mediate protection of newly-synthesized DNA during HU-induced replication stress. To address this issue, the serine or threonine residues at each of the seven S/TQ sites (residues 1308, 1316, 1330, 1351, 1386, 1417 and 1569) were mutated either to alanine to abolish phosphorylation (*rif1-7A*), or else to glutamic acid to mimic phosphorylation (*rif1-7E*) (Fig 4A). To confirm that *rif1-7A* and *rif1-7E* were expressed at similar levels to wild type Rif1, a 13xMyc tag was introduced at the C-termini of the mutant alleles. Western blot analysis of G1 phase-blocked cells carrying the *rif1-7A* and *rif1-7E* alleles showed similar expression levels to a strain with similarly tagged wild type Rif1 (Fig 4B).

Effects on nascent DNA protection were then tested using the procedure in Fig 1A. For the *rif1-7E* mutant, we found a slight but significant reduction in the extent of initial progression of replication forks during the initial IdU labelling period (e.g. median length 12.4 kb in wild-type versus 11.2 kb in the *rif1-7E* mutants, Fig. 4C), the reason for which is unclear. With regard to nascent DNA stability after HU blockage, we found that the *rif1-7A* non-phosphorylatable allele shows a significant decrease in nascent DNA tract length (from 12.4 kb to 8.7 kb after 1 hour of HU block, Fig 4C), a reduction comparable to the decrease in nascent DNA tract length in *rif1Δ* (which in this experiment showed a reduction in tract length from 12.4 kb to 8.7 kb over the same interval, Fig 4B). A repeat of the experiment produced very similar results (Fig S2). This defect in nascent DNA protection in *rif1-7A* suggests that phosphorylation of the S/TQ site cluster may be important for nascent DNA protection. Consistent with this idea, the phosphomimic *rif1-7E* allele in contrast showed no defect in the protection of DNA in a HU block, with the initial tract length of 11.2 kb maintained at 12.5 and 11.2 kb at 1 and 1.5 hr after HU addition (Fig 4B, also see Fig S2). This result is consistent with the *rif1-7E* allele being constitutively competent for nascent DNA protection. We conclude that, in order for Rif1 to protect nascent DNA, one or more of the clustered S/TQ checkpoint sites must be phosphorylated by the Tel1/Mec1 PIKK kinases. It therefore appears that Rif1 is an important target for phosphorylation after HU-induced replication stress, to enable nascent DNA protection.

## Discussion

Here we have shown that *S. cerevisiae* Rif1 protects newly synthesized DNA at HU-induced stalled forks by interacting with PP1/Glc7. We found that in the absence of Rif1 the Sgs1-Dna2 helicase-nuclease complex is primarily responsible for degrading nascent DNA (Fig 4D). Our results reveal the mechanism through which Rif1 protects nascent DNA is conserved from budding yeast to humans.

We discovered moreover that the S/TQ phospho-site cluster located within the unstructured region of *S. cerevisiae* Rif1 is required for nascent DNA protection. Specifically, a non-phosphorylatable *rif1-7A* allele caused a nascent DNA protection defect comparable to that of a full *rif1Δ* deletion. A phosphomimic *rif1-7E* allele in contrast did not produce any defect, supporting the suggestion that checkpoint-mediated phosphorylation of the Rif1 S/TQ cluster is crucial to protect nascent DNA at stalled forks. The function of the Rif1 S/TQ cluster has been the subject of debate, especially since mutating sites in this cluster does not impact telomere length in an otherwise wild-type background (although some effect on telomeres was observed in the context of *rif2* or *tel1* mutations) [48]. Our results now assign a clear physiological function for the ST/Q site cluster phosphorylation as being important for nascent DNA protection.

How Rif1 is recruited to stalled forks is still mysterious, and one interesting possibility is that Mec1/Tel1-mediated phosphorylation stimulates the association of Rif1 with replication forks upon checkpoint activation. Our results show that phosphorylation of the potential Tel1/Mec1 S/TQ sites in the Rif1 unstructured region is crucial for its function in protecting nascent DNA. Phosphorylation of these sites in response to activation of the intra-S phase checkpoint pathway may act as a signal to recruit Rif1 to stalled forks (Fig 4D). In human HeLa cell lines and *Drosophila*, Rif1 has been shown to interact with progressing replisomes. In *Drosophila*, fork association is largely dependent on Suppressor of Underreplication protein (SUUR) [51,52]. Rif1 also appears to be recruited to stalled replication forks in mouse embryonic fibroblasts (MEFs) [18]. In budding yeast, ChIP profiles suggest that Rif1 is recruited to stalled forks [30], but not to unperturbed forks. The fact that a Rif1-ΔC594 mutant, which lacks most of the ST/Q site cluster, is defective for recruitment to stalled forks [30] supports the notion that checkpoint phosphorylation is important for stalled fork recruitment of Rif1. Especially since the SUUR protein is not conserved in either yeast or human cells, at this point the interaction partners important for recruiting yeast and human Rif1 to stalled replication forks are unclear. Our results raise the possibility that the recruitment mechanism may involve recognition of RIF1 checkpoint phospho-sites by an unidentified replisome factor. Alternatively, phosphorylation may cause allosteric changes in Rif1 structure, which may in turn allow an association with components in replication forks. Replication stress also stimulates phosphorylation of RIF1 at a cluster of sites in human cells (Hiraga, Watts, & Donaldson, unpublished), so aspects of this recruitment mechanism could be conserved.

Our results also show that PP1 is required for Rif1 to mediate nascent DNA protection. The target dephosphorylated by this regulation is not completely clear, but the Dna2-WRN/Sgs1 complex is a good candidate, given its primary responsibility for nascent DNA degradation in both yeast and mammals lacking RIF1 (Fig 4D). Previous work in mammalian cells showed that both DNA2 and WRN (one of two mammalian homologs of yeast Sgs1) are hyperphosphorylated in the absence of RIF1 [17,18], suggesting that either or both DNA2 and WRN may indeed be direct targets of RIF1-PP1. MEFs treated with a PP1 inhibitor show hyperphosphorylation of DNA2 as assessed by Western blotting [18]. In Rif1-depleted human (HEK293-derived) cells, mass spectrophotometry analysis identified hyperphosphorylation of several residues in the WRN helicase either in untreated or HU-blocked conditions [17].

In yeast, phosphorylation of *S. cerevisiae* Sgs1 has been proposed to be involved in checkpoint activation through enhancing RPA and Rad53 interaction [53], but any effect of Sgs1 phosphorylation on nascent DNA stability has not been investigated. In *S. pombe*, checkpoint-mediated phosphorylation of Dna2-S220 has been proposed to enable Dna2 recruitment to replication forks and the formation and cleavage of regressed forks [54]; however, whether any equivalent phospho-site is important in *S. cerevisiae* to control events at blocked forks has not been addressed. In budding yeast, Dna2 is phosphorylated by the Cdk1 and Mec1 kinases [55]. Cdk1 phosphorylates residues T4, S17 and S237 both *in vitro* and *in vivo*, and mutating these sites to alanine leads to less processive resection of DSBs [55]. However there has been no investigation of how phosphorylation affects the activity of *S. cerevisiae* Dna2-Sgs1 at blocked forks. Nonetheless, since phosphorylation has been suggested to activate helicase and nuclease activities of Dna2-Sgs1, Rif1-PP1 could potentially counteract these activities by removing activating phosphorylations. A detailed, systematic study on the effect of phosphorylation on the combined Dna2-Sgs1 functional activities will need to be fully completed to understand how Rif1-PP1 may impact the function of this complex in nascent DNA protection.

While Sgs1 and Dna2 are good candidates, the possibility remains that Rif1-PP1 dephosphorylates other substrates to limit nascent DNA tract degradation. Various other components affect DNA protection at HU-stalled forks. For example the MRX complex acts in concert with chromatin modifiers including Set1 (catalytic component of the COMPASS complex that carries out H3K4 methylation) for remodelling of nascent chromatin to allow access by downstream helicases/nucleases to progressively resect DNA ends [26]. Another COMPASS component, Bre2, shows increased phosphorylation in a *rif1Δ* mutant (data not shown), highlighting that Rif1 could potentially affect nascent DNA protection through COMPASS or other complexes [56].

To summarise, we have found that *S. cerevisiae* Rif1 protects nascent DNA by acting with PP1 to oppose the Dna2-Sgs1 helicase-nuclease, in a mechanism conserved from yeast to human cells. In this context, it will be of particular interest to discover if checkpoint-mediated phosphorylation of human Rif1 is also important for protecting newly synthesized DNA during replication stress, and exactly why checkpoint phosphorylation is critical for this particular function of yeast Rif1.

## Materials and Methods

### Yeast strains

Yeast strains used for this study were all in a W303 *RAD5*^*+*^ background and are described in Table 1. Strains VGY85 and CMY6 were previously described [30,57]. CMY42 was generated in a two-step process. First, a region of the N-terminus of *RIF1* (bases 97-2508) was replaced by a *URA3* cassette. This *URA3* cassette was then replaced using a PCR fragment amplified from plasmid pSH192 [12] containing mutations of the PP1 binding sites of *RIF1*. CMY46, CMY47, CMY52 and CMY53 were created by replacing the *EXO1* or *SGS1* genes with *TRP1* or *URA3* respectively, by one-step PCR replacement. To construct CMY128, first a CRISPR Cas9 plasmid was made to enable introduction of the *dna2-1* mutation.

**Table 1.** Yeast strains used in this study.

| Name | Relevant genotype | Background | Source/<br>reference |
| --- | --- | --- | --- |
| YK402 | *MATa bar1Δ::hisG ade2-1 can1-100 his3-11,15 leu2-3,112 trp1-1 ura3-1* | W303 *RAD5<sup>+</sup>* | [12] |
| YSM20 | YK402 *Rif1-13Myc::HIS3MX6* | W303 *RAD5<sup>+</sup>* | [49] |
| VGY85 | YK402 *trp1-1Δ::BrdU-InC-KanMX4* | W303 *RAD5<sup>+</sup>* | [57] |
| CMY6 | VGY85 *rif1Δ::HIS3* | W303 *RAD5<sup>+</sup>* | [30] |
| CMY42 | VGY85 *rif1-pp1bs* | W303 *RAD5<sup>+</sup>* | This study |
| CMY46 | VGY85 *exo1Δ::TRP1* | W303 *RAD5<sup>+</sup>* | This study |
| CMY47 | CMY6 *exo1Δ::TRP1* | W303 *RAD5<sup>+</sup>* | This study |
| CMY52 | VGY85 *sgs1Δ::URA3* | W303 *RAD5<sup>+</sup>* | This study |
| CMY53 | CMY6 *sgs1Δ::URA3* | W303 *RAD5<sup>+</sup>* | This study |
| CMY54 | VGY85 *ura3-1::GALpr-OsTIR1 codon plus-URA3* | W303 *RAD5<sup>+</sup>* | This study |
| CMY56 | CMY6 *ura3-1::GALpr-OsTIR1 codon plus-URA3* | W303 *RAD5<sup>+</sup>* | This study |
| CMY58 | CMY54 *DNA2-AID::natmx* | W303 *RAD5<sup>+</sup>* | This study |
| CMY59 | CMY56 *DNA2-AID::natmx* | W303 *RAD5<sup>+</sup>* | This study |
| CMY71 | VGY85 *rif1Δscd::URA3* | W303 *RAD5<sup>+</sup>* | This study |
| CMY128 | VGY85 *dna2-1* | W303 *RAD5<sup>+</sup>* | This study |
| CMY130 | CMY71 *rif1-7A* | W303 *RAD5<sup>+</sup>* | This study |
| CMY132 | CMY71 *rif1-7E* | W303 *RAD5<sup>+</sup>* | This study |
| CMY134 | CMY128 *rif1Δ::HIS3* | W303 *RAD5<sup>+</sup>* | This study |
| CMY135 | CMY130 *rif1-7A-13xMyc::HIS3MX6* | W303 *RAD5<sup>+</sup>* | This study |
| CMY136 | CMY130 *rif1-7A-13xMyc::HIS3MX6* | W303 *RAD5<sup>+</sup>* | This study |
| CMY137 | CMY132 *rif1-7E-13xMyc::HIS3MX6* | W303 *RAD5<sup>+</sup>* | This study |
| CMY138 | CMY132 *rif1-7E-13xMyc::HIS3MX6* | W303 *RAD5<sup>+</sup>* | This study |
| CMY139 | CMY132 *rif1-7E-13xMyc::HIS3MX6* | W303 *RAD5<sup>+</sup>* | This study |

**Table 2.**
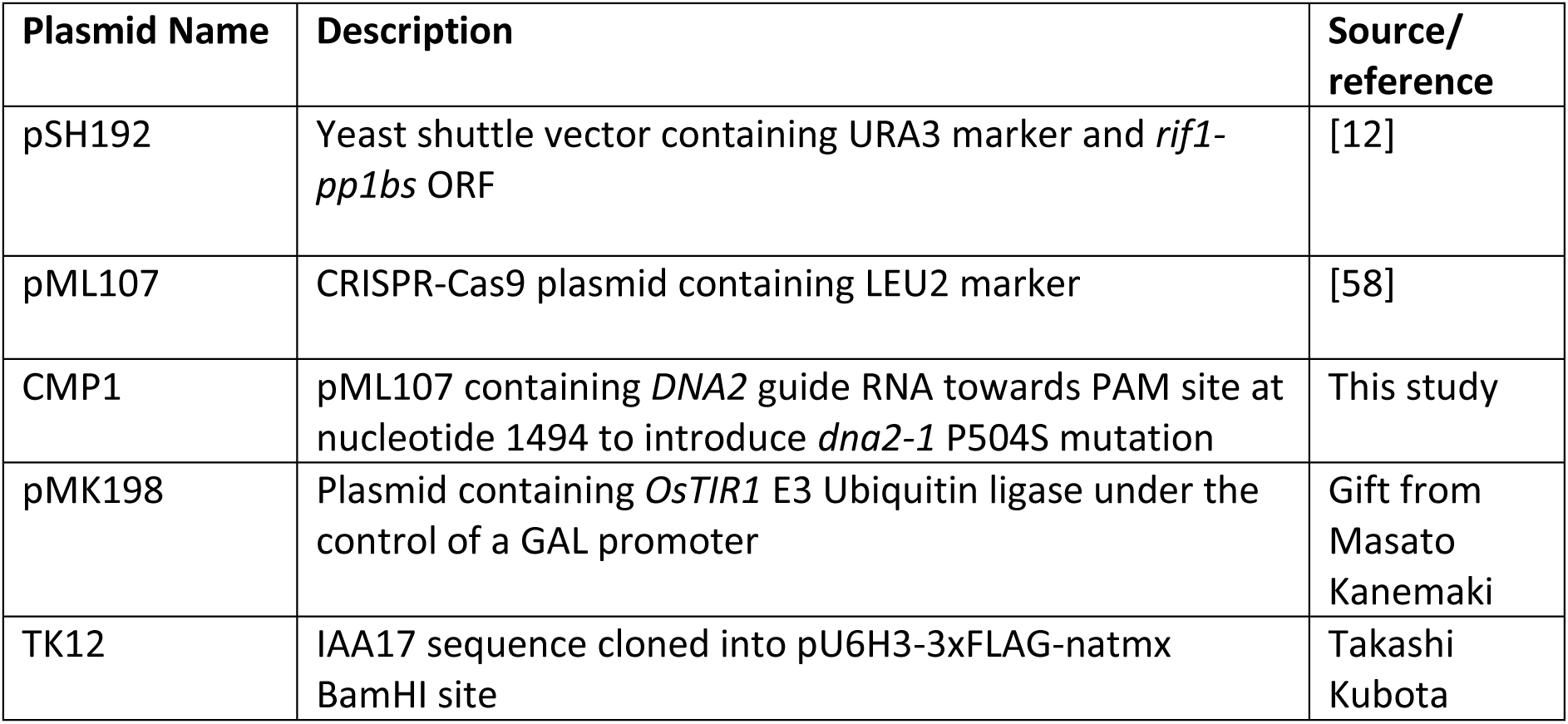
Plasmids used in this study.

**Table 3.**
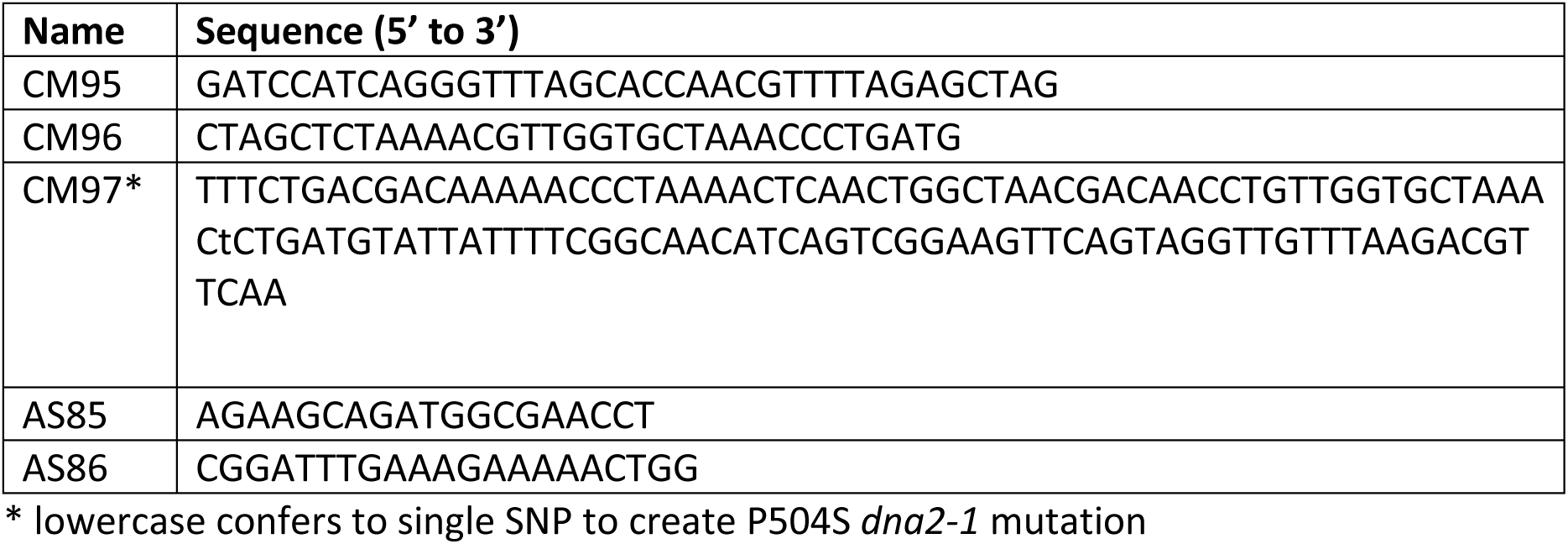
Primers used in this study.

CRISPR Cas9 plasmid pML107 [58] was digested with BclI and SwaI restriction enzymes. The primer pair CM95-CM96, which encode guide RNA directed towards *DNA2*, were annealed and cloned into the linearised plasmid to create plasmid CMP1. Primer CM97 was used as a ssDNA repair template to introduce the *dna2-1* mutation P504S. After transformation with CMP1 and the repair template, introduction of the correct mutation was confirmed by sequencing. *RIF1* was then replaced in CMY128 with a *HIS3* cassette, by one step PCR replacement, to create CMY134. Plasmid pMK198 (a gift from Masato Kanemaki), which contains the E3 ubiquitin ligase *OsTIR1* under the control of a *GAL* promoter, was digested with StuI and integrated into the genome of VGY85 and CMY6 at the *ura3-1* locus. The C-terminus of *DNA2* was tagged with full length AID amplified from plasmid TK12, which also included a 3xFLAG tag and nourseothricin selection marker to create CMY58 and CMY59.

Integration of the AID tag was confirmed by sequencing. Rif1 phospho-site mutants were constructed by first replacing the Rif1 ORF nucleotide sequence 3903-4724 (amino acids 1302-1574) with a *URA3* cassette, removing the entire cluster of seven S/TQ sites to create a *rif1Δscd::URA3* strain, CMY71. From IDT Technologies we obtained 822 bp ‘gBlock’ fragments encoding either alanine residues (7A allele) or glutamic acid residues (7E allele) instead of serine/threonine at the seven S/TQ sites. These phospho-site mutant fragments were transformed into CMY71 to replace the *URA3* cassette, creating the *rif1-7A* or *rif1-7E* strains CMY130 and CMY132, respectively. The S/TQ site mutations were confirmed by sequencing. The C-termini of the *rif1-7A* and *rif1-7E* alleles were tagged with a Myc tag by amplifying a 13xMyc-HIS3MX6 casette from YSM20 [49] genomic DNA using primers AS85-AS86, and transforming the amplified fragment into CMY130 and CMY132. Creation of these *rif1-7A* and *rif1-7E* Myc-tagged alleles was confirmed by sequencing.

## DNA combing

### DNA combing was performed as previously described [30]. Briefly, cells were arrested with

α-factor, then collected by centrifugation and resuspended in fresh media containing 1.4U /litre Pronase (to release cells into S phase) and 1.13 mM IdU (to label nascent DNA) and cultivated at 30°C. Cells were collected by filtration, washed and resuspended in fresh media containing 0.2 M HU and 5 mM thymidine. Thymidine was included to minimise labelling of ongoing DNA synthesis by any residual IdU. Cells were collected after 0, 1 and 1.5 hr and encased in low melting agarose plugs. Cells in plugs were spheroplasted and genomic DNA prepared using FiberPrep DNA extraction kit (Genomic Vision), according to manufacturer’s instructions. DNA combing was performed using FiberComb Instrument (Genomic Vision). Coverslips with combed DNA were probed with anti-IdU (Becton Dickinson 347580) and anti-ssDNA (Millipore MAB3034) followed by appropriate secondary antibodies with fluorescent conjugates for immunodetection. IdU tracts were visualised under a Zeiss Axio Imager.M2 microscope equipped with Zeiss MRm digital camera with a Zeiss Plan-Apochromat 63x/1.40 Oil objective lens. Images were analysed using ImageJ software. IdU-labeled tract lengths were measured using the following criteria: tracts must be at least 2 µm in length; be separated from each other by 5 µm or more; lie on a ssDNA fragment at least 50 µm in length with the tract finishing at least 5 µm from the end as visualised by ssDNA antibody. IdU tract length (in µm) was converted to kilobases using the predetermined value (2 kb/µm) for the DNA combing method.

### *dna2-1* growth plate assay

To verify temperature sensitivity of *dna2-1* mutants, strains were grown overnight in YPD. 2.5×10^5^ cells/ml were collected and serially diluted 1:5 onto YPD plates and incubated at 23°C or 30°C.

### Dna2 depletion

To investigate the effect of Dna2-AID depletion on cell viability, cells were grown overnight in YPD and 1×10^7^ cells/ml serially diluted 1:10 the next day onto YPD, or YP+2% galactose and where required supplemented with auxin (final concentration 1 mM).

For Dna2 depletion in liquid culture using the auxin degron system, cells were grown overnight in YP+2% raffinose and arrested in G1 phase using α-factor for 2 hours. Galactose was added to a final concentration of 2% to induce expression of the E3 ubiquitin ligase OsTir1. After 1 hour, auxin (final concentration 1 mM) was added to deplete Dna2. For experiments involving labelling of nascent DNA, Dna2-depleted cells were pre-incubated with 1.13 mM IdU for 15 minutes. 1.4U /litre Pronase was then added directly (without filtration) to allow release into S phase with nascent DNA labelling. Media was filtered and cells were resuspended in YEP 2% galactose, 1 mM auxin, 0.2 M HU and 5 mM thymidine to initiate the HU block. Samples were taken after 0, 1 and 1.5 hours and DNA combing performed as previously described [30].

### Western Blotting

To measure Dna2-AID degradation, cells were arrested with α-factor as outlined above and a ‘-60 min’ sample was collected. Galactose was added as above to a final concentration of 2%. 1 hr later auxin (final concentration 1 mM) was added and samples collected after 0, 15, 30 and 60 minutes. Proteins were prepared using the alkaline extraction method [59]. 175 µg and 5 µg of samples were loaded onto mini-PROTEAN 4-15% TGX gels (BIORAD) for western blotting and SYPRO staining, respectively. Dna2-AID 3xFLAG and loading control tubulin were detected using anti-FLAG M2 antibody (Sigma, F1804) and α-Tubulin YOL 1/34 (Santa Cruz), respectively.

To assess *rif1-7A* and *rif1-7E* expression levels, cells were arrested with α-factor for 2 hours and samples collected. Proteins were prepared as above. 20 µg of samples were loaded onto a mini-PROTEAN 4-15% TGX Stain-Free gel (BIORAD) for western blotting. Rif1-Myc was detected using anti-Myc antibody (MBL 047-03) and loading control tubulin detected as mentioned above.

## Acknowledgments

We thank Masato Kanemaki for constructs. This work was supported by Cancer Research UK Programme Award C1445/A19059 to AD and SH.

## Supporting Information

**Fig S1.**
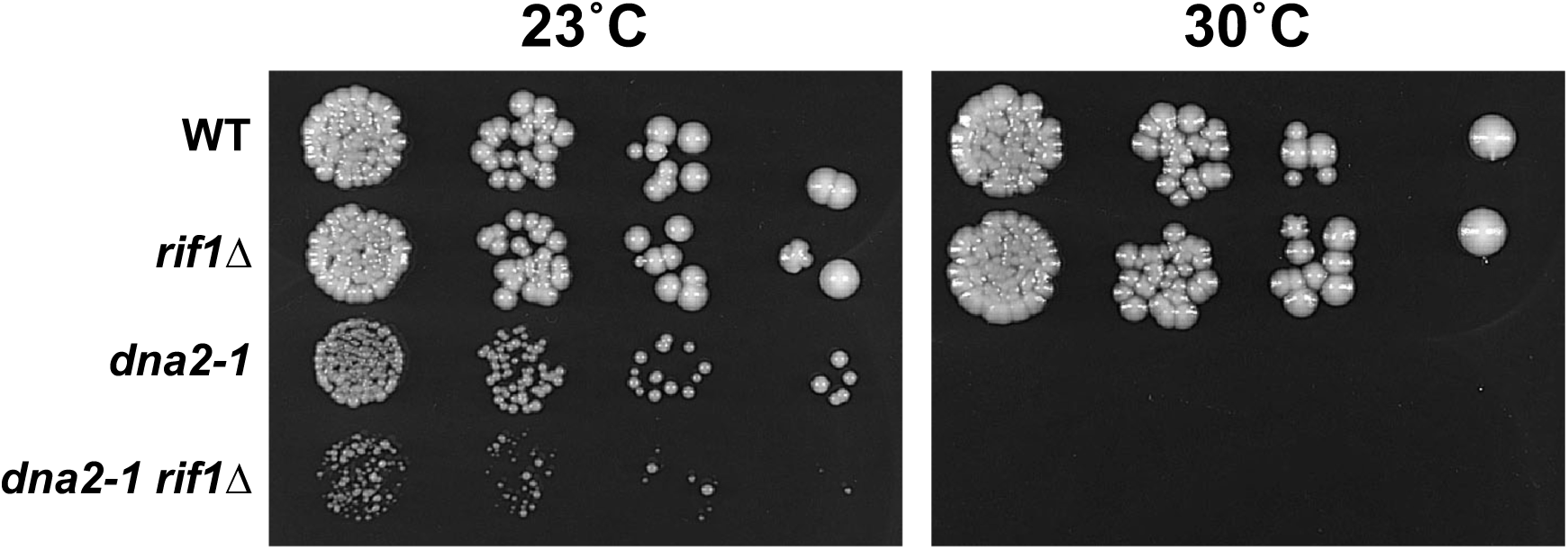
Deletion of Rif1 in a *dna2-1* background leads to synthetic sickness. Serial dilutions (1:5) of cells grown on YPD at 23°C and 30°C. *dna2-1* mutants are temperature sensitive and fail to grow above 30°C. Plates were imaged after 4 and 3 days respectively.

**Fig S2.**
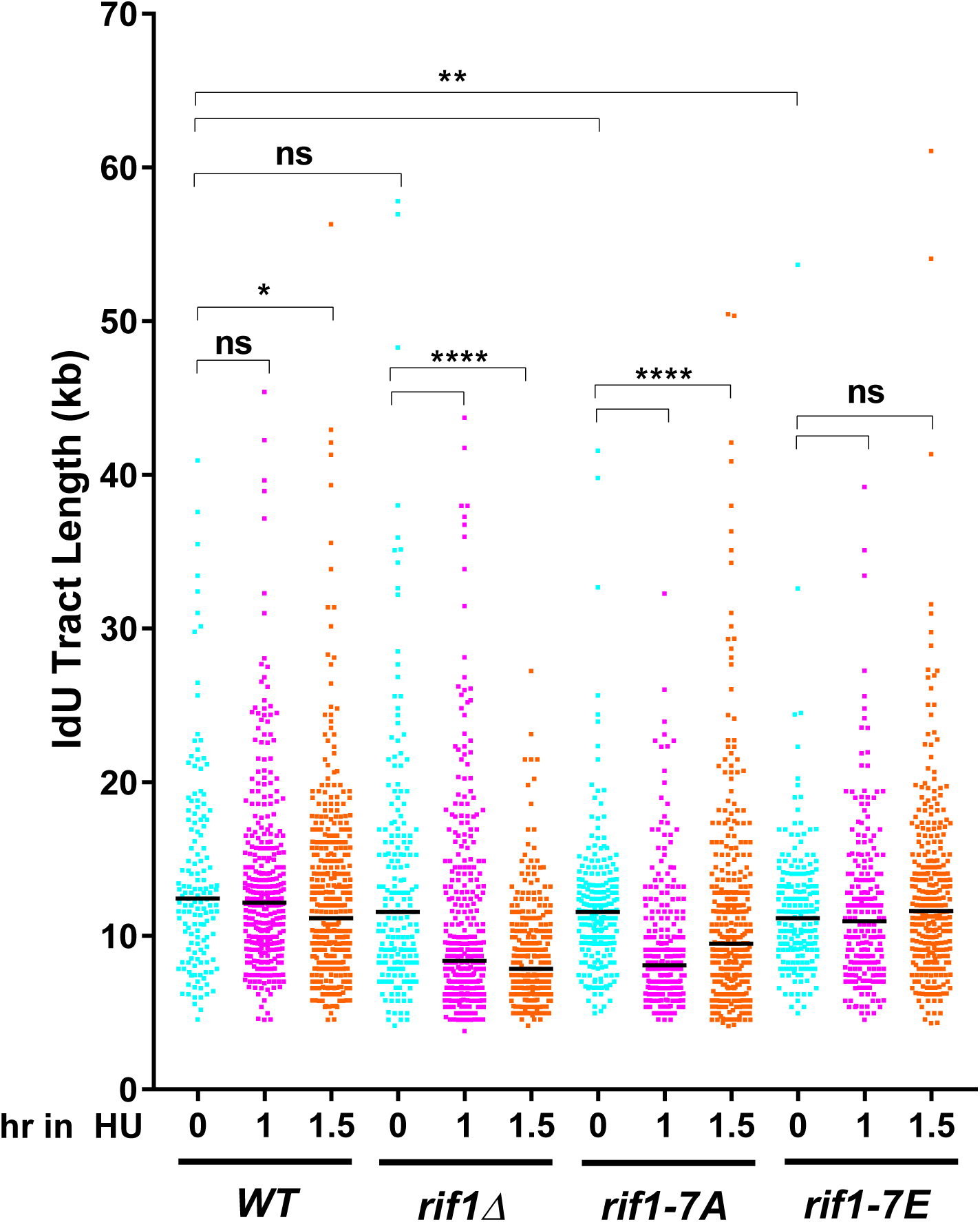
Nascent DNA protection requires a cluster of Tel1/Mec1 checkpoint recognition sites in the unstructured Rif1 C-terminal domain (repeat of experiment shown in Fig 4C). Analysis of the Rif1 S/TQ phospho-dead mutant *rif1-7A* reveals a defect in the protection of nascent DNA during a HU block, while fork protection is intact in the phosphomimic mutant *rif1-7E*. Black horizontal bars indicate median values. *, ** and **** indicates *p*-values less than 0.05, 0.01 and 0.0001, respectively. Obtained by Mann-Whitney-Wilcoxon test. Ns means “not significant”.

**Table S1. Compiled nascent IdU tract measurements**

